# Membrane-binding domains define REMORIN phylogeny and provide a predicted structural basis for distinctive membrane nano-environments

**DOI:** 10.64898/2025.12.22.695504

**Authors:** David Biermann, Julien Gronnier

## Abstract

REMORIN (REM) proteins are structural components of the plant plasma membrane that modulate membrane nano-organization and biophysics. They have been proposed to function as versatile scaffolds in the context of hormone signaling, immunity and symbiosis. REMs have been classified into six groups based the length and the amino acid composition of their intrinsically disordered N-terminal domain. Here we show that REM phylogeny is dominated by the evolution of their conserved C-terminal domain and defines four major REM clades. Structural bioinformatics analyses predict the conservation of a putative membrane binding interface formed by REM C-terminal domains and reveal a striking diversity in their curvatures and lengths. A subset of REMs is predicted to form C-terminal domain-mediated higher-order oligomers providing an additional level of diversity in REM membrane-binding interfaces. We discuss the implications of the predicted variations in REM C-terminal domain structure for their molecular function and membrane organization.

## Introduction

REMORINs (REMs) constitute a plant-specific protein family characterized by a conserved and structured C-terminal domain and divergent and intrinsically disordered N-terminal domain (Marín & Ott, 2012; Perraki et al., 2012; Raffaele et al., 2007) REMs have been classified into six groups based on the length and the amino acid composition of their N-terminal domains (Raffaele et al., 2007). In recent years, REMs have emerged as regulatory components of various physiological processes (Gouguet et al., 2021) and are best characterized for their role plant-microbe interactions. For instance, group 1 REMs regulate cell-to-cell movement of viruses belonging to diverse genera (Cheng et al., 2020; Fu et al., 2018; Huang et al., 2019; Jolivet et al., 2025; Perraki et al., 2014; Perraki et al., 2012; Perraki et al., 2018; Raffaele et al., 2009; Rocher et al., 2022). Upon perception of bacterial-derived microbe-associated molecular patterns two *Arabidopsis thaliana* REM paralogues, AtREM1.2 and AtREM1.3, form nanodomains that contain the actin nucleating protein FORMIN 6 and modulate actin nucleation (Ma et al., 2022) Similarly, salt sensing induces AtREM1.2 and FORMIN 6 co-clustering (von Arx et al., 2025). In leguminous species, the symbiotic REM1 (SYMREM1) has been linked to receptor kinases (Lefebvre et al., 2010; Liang et al., 2018), and shown to regulate membrane topology (Su et al., 2023) during the establishment of nodule symbiosis. REMs have served as model proteins to study the mechanisms regulating plant plasma membrane nanoscale organization and dynamics (Gronnier et al., 2017; Jarsch et al., 2014; Liang et al., 2018). The plasma membrane organization and dynamics have been predominantly studied for group 1 REMs. Long-term single molecule imaging showed that Arabidopsis REM1.2 is dynamic and transiently spatially arrested in confined membrane nano-environments (von Arx et al., 2024). Their nanoscale organization is responsive to stimuli (Ma et al., 2022)(von Arx et al., 2025), modulated by post-translational modifications (Fu et al., 2018; Jolivet et al., 2025; Konrad et al., 2014; Perraki et al., 2018) as well as by molecular partners (Huang et al., 2019; Jolivet et al., 2025; Jolivet et al., 2023; Liang et al., 2018).

The plasma membrane association and nanoscale dynamics of REMs are regulated by direct interactions with anionic phospholipids (Gronnier et al., 2017; Legrand et al., 2023; Legrand et al., 2019; Perraki et al., 2012; Xu et al., 2024). REMs associate with the plasma membrane inner leaflet via their C-terminal domain which is predicted to assemble into anti-parallel dimers forming a membrane binding interface (Su et al., 2023; Xu et al., 2024) which includes the characterized lipid-binding REM C-terminal anchor (Gronnier et al., 2017; Konrad et al., 2014; Perraki et al., 2012). REMs influence membrane biophysical properties (Gronnier et al., 2017; Huang et al., 2019; Legrand et al., 2019) and cell membrane shape (Su et al., 2023). While a few REMs have been studied in detail *in vitro* and *in vivo* for their plasma membrane organization (Gronnier et al., 2017; Legrand et al., 2019; Ma et al., 2022; Su et al., 2023) and have been used as plasma membrane markers for cell biological studies (Bücherl et al., 2017; Cui et al., 2024) relatively limited genetic characterization is limiting our understanding of the cellular and molecular processes regulated by REMs. This can be explained in part by their genetic redundancy and by the absence of a bona fide phylogenetic classification of the REMORIN protein family.

Here, we performed a phylogenetic analysis of REM protein family. We found that amino acid variation in REM C-terminal domain classifies REMs into four major clades, which substantially revise relationships between REM homologues. Using structural bioinformatics, we observed the predicted conservation of a REM C-terminal domain-mediated membrane binding interface, as well as the diversity in their predicted curvature and length, and their intrinsic capacity to mediate the higher-order oligomers. We discuss the potential implication of these predictions for the structuration of the plasma membrane at the nanoscale.

## Results

### A phylogenetic classification of REMORIN protein family

The classification of REM in six distinct groups has been predominantly defined based on variation in the length and amino acid composition of their N-terminal domains (Raffaele et al., 2007), a classification which does not align with REM phylogeny (Bozkurt et al., 2014; Cai et al., 2020; Checker & Khurana, 2013; Raffaele et al., 2007; Toth et al., 2012; Zhang et al., 2020). We therefore aimed to clarify REMs phylogenetic relationship. We use the protein sequences of twenty REMs, the seventeen encoded in *Arabidopsis thaliana* genome (Col-0) and the three REMs encoded in the genome of *Marchantia polymorpha* (Tak-1), to perform iterative BLAST searches in 53 species covering the green lineage and collected more than 1100 REM sequences (Supplementary data 1). We constructed consensus phylogenetic trees based on REM full length sequences and on the conserved C-terminal domain. Both phylogenetic analyses converge in describing the classification of REMs into four major clades with strong branch support indicating that REM phylogeny is dominated by the evolution of the REM C-term domain. These analyses substantially revise relationships between REM homologues (Fig. 1). Group 6 REMs are distributed in two different clades (II and IV). The Arabidopsis group 3 REMs, AtREM3.1 and AtREM3.2, belong to distinct clades (I and II) and likely correspond to truncated duplications of group 6 and group 1 REMs, respectively (Fig. 1). The group 5 REMs are found in clade IV together with group 6 REMs. Finally, group 2 REMs (e.g., MtREM2.2, Lefebvre et al., 2010) cluster together with group 1 REMs in clade I. Amino acids sequence variations of the REM C-term domain further subdivide clade III into two subclades (IIIa and IIIb) and the clade IV into three subclades (IVa, IVb and IVc). Amino acids sequence variations of the REM N-terminal domains divide clade II into two sub-clades (IIa and IIb) and the subclade IVc into two groups (IVc-1 and IVc-2). We found REM representatives in non-seed plants (bryophytes, and ferns) related to clades I, III and IV, suggesting that these clades stem from ancient duplication events and may be linked to early functional diversification (Fig. 1). Clade II is constituted by REMs only found in angiosperm spermatophytes and may represent recent functional diversification potentially linked to the emergence of seed plants. Alternatively, clade II REMs may have been lost in non-seed plants. Altogether, these phylogenetic analyses revisit REM classification and provide a basis for the genetic characterization of REM in model plants and crop species alike.

**Figure 1.**
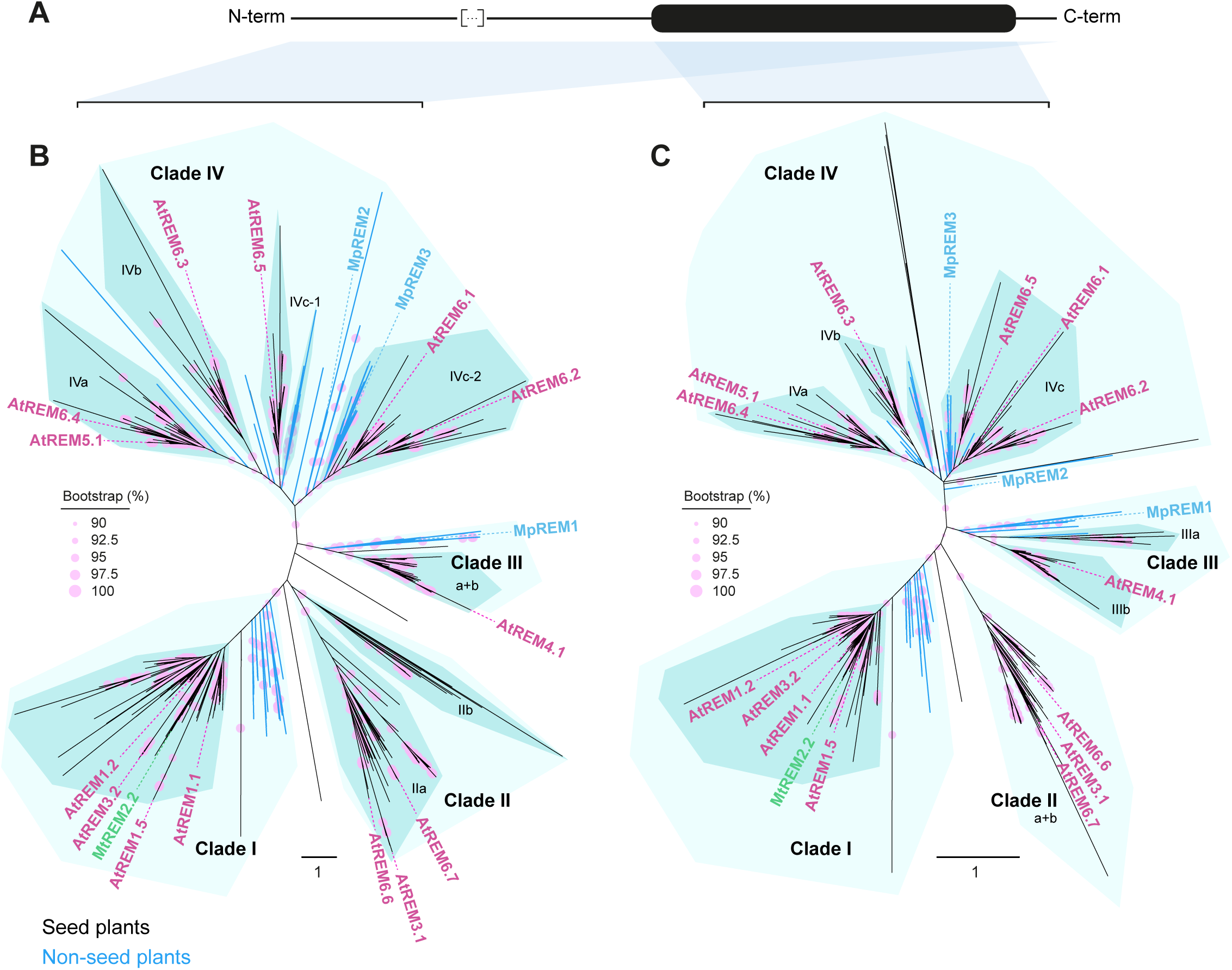
REMORIN phylogeny is dominated by C-terminal domain evolution. **A**. Schematic representation of the canonical REMORIN protein secondary structure depicting an alpha helix with a thick line and coil region with a thin line. The shade areas highlight REMORIN segments used to construct the phylogenetic trees presented in B and C. **B-C**. Maximum likelihood trees of 1133 full length (**B**) and C-terminal domain alpha helix (**C**) REMORIN proteins. Total tree length equal 640.776 (**B**) and 208.659 (**C**), and Log-likelihood equal −377021.439 (**B**) and −70993.460 (**C**). Branches support is indicated in % from 2000 bootstraps per tree. The scale bar indicate substitution per site. The position of representative REMORIN members *Arabidopsis thaliana* (At) REM1.1 (AT3G48940), AtREM1.2 (AT3G61260), AtREM1.5 (AT1G63295), AtREM3.1 (AT1G69325), AtREM3.2 (AT4G00670), AtREM4.1 (AT3G57540), AtREM5.1 (AT1G45207), AtREM6.1 (AT2G02170), AtREM6.2 (AT1G30320), AtREM6.3 (AT1G53860), AtREM6.4 (AT4G36970), AtREM6.5 (AT1G67590), AtREM6.6 (AT1G13920), AtREM6.6 (AT5G61280), *Medicago truncatula* REM2.2 (Medtr8g097320) and *Marchantia polymorpha* (Mp) REM1 (Mp2g17160.1), MpREM2 (Mp1g28990.1) and MpREM3 (Mp2g06670.1) is indicated.

### REMORIN C-terminal domains provide a predicted structural basis for distinctive membrane nano-environments

We observed that REM phylogeny is primarily dominated by amino acids sequence variations within their C-terminal domains (Fig. 1). In line with previous observations, REM C-terminal domains show a higher degree of conservation in all phylogenetic clades (Fig. S1). *In silico* modeling of *Arabidopsis thaliana* and *Medicago truncatula* REMs using AlphaFold (Jumper et al., 2021) predicted the formation of C-terminal domain-mediated REM anti-parallel dimers (Su et al., 2023; Xu et al., 2024). In agreement, size exclusion chromatography of recombinant His-SYMREM1 showed the formation of homodimers *in vitro* (Su et al., 2023) and *in silico* molecular dynamics simulations further highlighted the stability of predicted anti-parallel dimers (Xu et al., 2024). These predicted structures are distinctively positively charged on their convex side, suggestive of a large membrane binding interface (Su et al., 2023; Xu et al., 2024). To infer the generality of these observations, we used AlphaFold3 (AF3), which offers substantially improved accuracy for the prediction for protein complexes (Abramson et al., 2024), to model the structure of an extended and diverse set of REMs. We randomly selected 43 REM representatives covering all four clades of the phylogenetic tree (Supplementary Table 2). In agreement with previous models (Su et al., 2023; Xu et al., 2024), all 43 REM C-terminal domains were predicted to form anti-parallel dimers (Fig. S2) indicating that REM dimerization is an evolutionary conserved feature. We observed that residues engaged in intermolecular interactions exhibit a high degree of conservation across REM protein family (Fig. S2), suggesting that these residues are under selective pressure and that REM dimerization is important for REM function.

Interestingly, we noticed variations in the curvature and the length of the putative membrane-binding interfaces (Fig.2; Fig. S2). We systematically estimated the curvature of REM dimers (Fig. S3) and observed variations that span one order of magnitude (Fig. 2B). Similarly, the length of predicted membrane binding interfaces varied from approximately 10 to 25 nm (Fig. 2C). We observed quantitative differences in these parameters between clades but no strict clade specificity (Fig. S4). Such variations could be found at the scale of a single plant species as observed for *Arabidopsis thaliana* (Fig. S5A). Structural modelling predicted that SYMREM1 and StREM1.3 could form higher order oligomers (Su et al., 2023; Xu et al., 2024). Through quantitative analyses of the AF3 predictions of Arabidopsis thaliana REM1.2 (AtREM1.2) homo-oligomers we observed that hexamer and octamers are predicted with the highest scores (Fig. 3A). Similarly, the clade III Lupinus albus Lalb_Chr20g0110781 REM, the clade I AtREM1.1 and the clade II AtREM4.1 were best predicted as hexamers and octamers (Fig. S5 and S6). AtREM1.2 hexamers and octamers exhibit a large area of positively charged residues decorating predicted membrane binding interfaces (Fig. 3B). Dimers, hexamers and octamers differ in the area and curvature of their predicted membrane binding interface (Fig. 3B). Interestingly, we observed that not all REMs are predicted to form higher-order homo-oligomers (Fig. S5 and S6), suggesting the existence of structural features specifying REM C-terminal domain-mediated oligomerization. Altogether, these observations unveil an unsuspected diversity in the predicted dimer- and oligomer-based membrane binding interfaces formed by REM protein family members.

**Figure 2.**
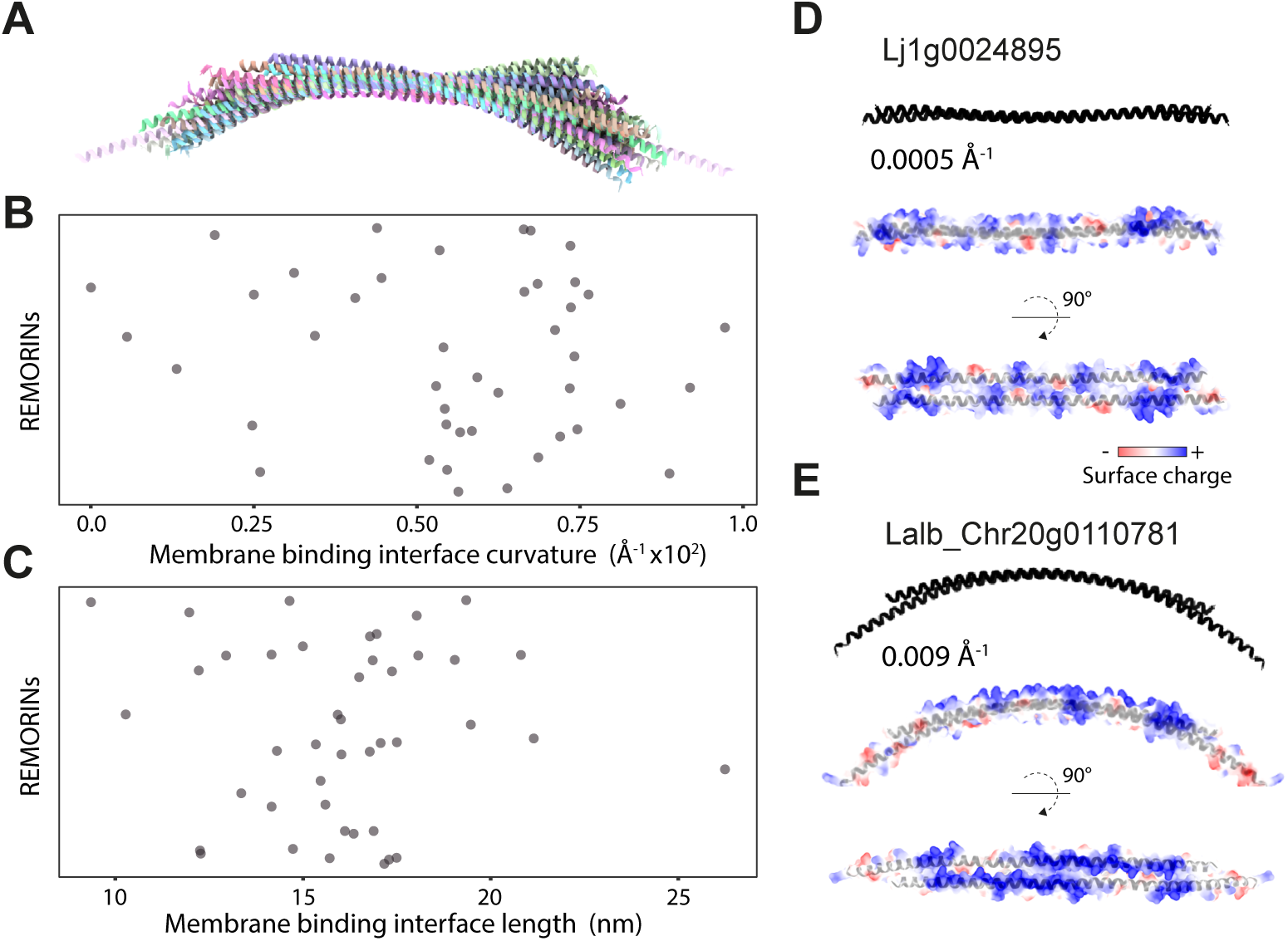
Structural analysis of predicted REMORIN membrane-binding domains. **A**. Superimposed AlphaFold3 structure predictions of 43 REMORIN C-terminal domain dimers. **B**. Curvatures of REM C-terminal domain dimers estimated using from nonlinear least-squares minimization circle fitting to AlphaFold3 predicted structures. See also Fig. S2. **C**. Length of REM membrane-binding interfaces estimated from the arc length of the fitted circle spanned by REM dimer Cα atoms. See also Fig. S2. **D-E**. AlphaFold3 predicted structures of *Lotus japanicus* (Lj) Lj1g0024895 (**D**) and *Lupinus albus* (La) Lalb_Chr20g0110781 (**E**) REM C-terminal domain dimer. The estimated curvater (Å^-1^) surface charges are represented.

**Figure 3.**
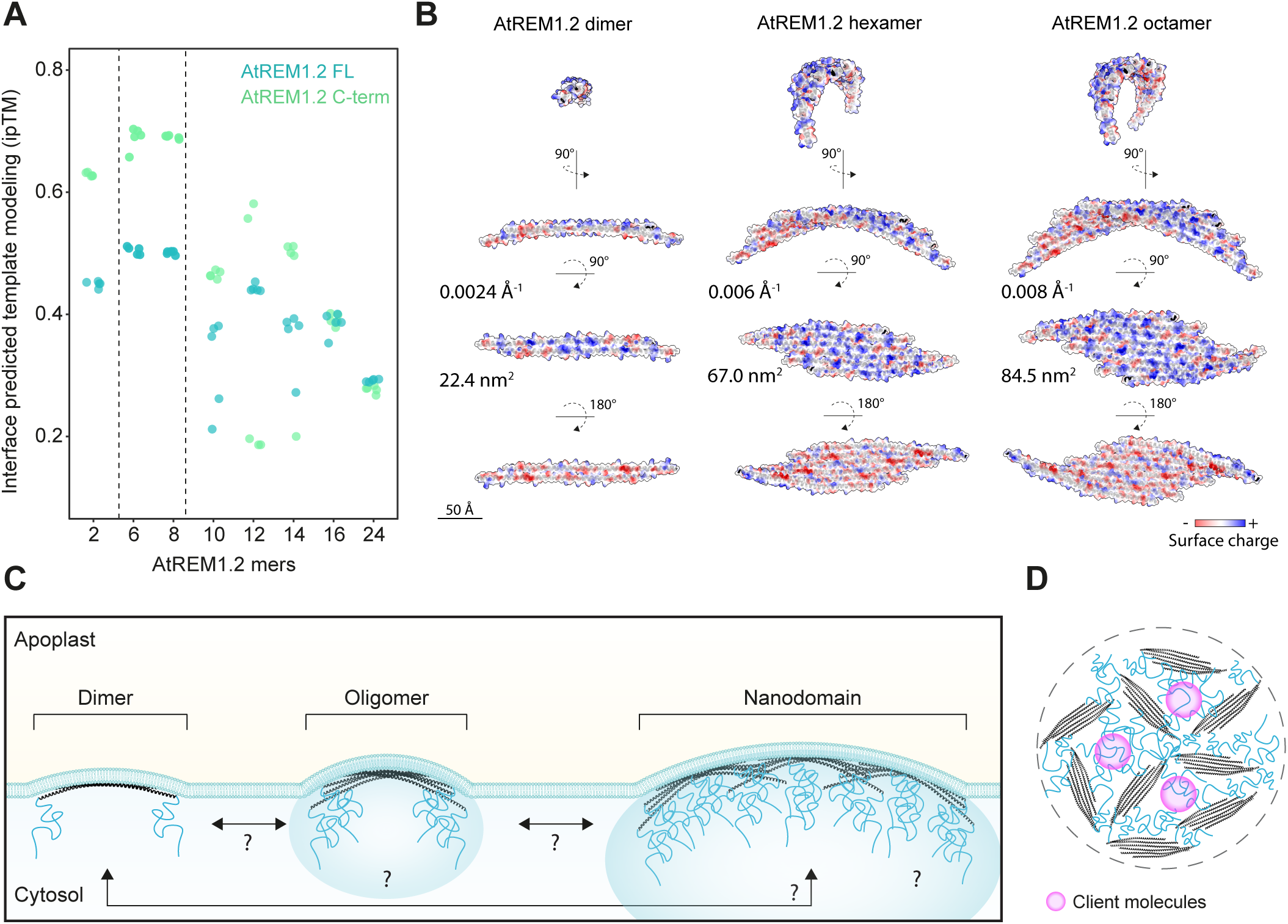
Quantitative analysis of predicted AtREM1.2 oligomerization status. **A**. Interface predicted template modelling (ipTM) score of AlphaFold3 predictions for full-length (FL) or C-terminal (C-Term) domain AtREM1.2 oligomers. Each datapoint represents individual ipTM prediction scores for each multimer. **B**. Predicted structures of AtREM1.2 dimer, hexamer and octamer. The estimated curvater (Å^-1^), the predicted membrane binding interface (nm^2^) and surface charges are represented. **C**. Schematic representation of the putative REM-membrane curvature relationship, REM dimers, oligomers and nanodomain may sense and/or impose various positive membrane curvatures. The strength or membrane curvatures is predicted to vary based on the intrinsic structural variations between REMs homologues. REM-membrane relationship can further be regulation by REM S-acylation and potential molecular condensate-like behavior of REM-oligomer- and REM-nanodomain-based molecular assemblies and the recruitment of REM-nanodomain client molecules (**D**). REM intrinsically disordered N-terminal domain is depicted in blue, and REM alpha helical C-terminal domain is depicted in black.

## Discussion

Our study clarifies REM protein family phylogeny. The classification of REMs into four major clades is not meant to replace the nomenclature for REM proteins (Raffaele et al., 2007), which has been widely used for nearly twenty years. Instead, it provides a basis for further genetic experimentation. REM phylogeny is dominated by variations in their conserved C-terminal domains. Our in-silico analyses, together with previous biophysical (Legrand et al., 2023; Xu et al., 2024) and cell biological studies (Gronnier et al., 2017; Ma et al., 2022; Su et al., 2023), indicate that REMs form diverse molecular assemblies, including dimers, oligomers and higher-order structures corresponding to REM nanodomains (Fig. 3C).

Antiparallel dimer formation is a predicted conserved feature of all tested REM (Fig. S2;(Su et al., 2023; Xu et al., 2024). REM dimers remain stable in molecular-dynamic simulations (Xu et al., 2024), and dimer-forming residues are under strong selective pressure (Fig. S2). These observations suggest that dimerization is core structural feature essential for REM function.

REM dimers are predicted to form membrane-binding domains with striking variation in curvature and length (Fig. 2). Curvature spans an order of magnitude, from near neutral (e.g., clade II REM Lj1g0024895) to strong positive curvature (e.g., clade III REM LalbChr20g0110781) potentially corresponding to strong convex membrane bending towards the extracellular space. Predicted REM C-terminal domain-mediated oligomerization is expected to provide additional variations in curvature (Fig. 3B-C), which could be further amplified by REM nanodomain formation (Fig. 3B).

Not all REMs are predicted to form REM C-terminal-domain-mediated homo-oligomers (Fig. S5 and S6). Among them, AtREM3.2, AtREM6.1 and AtREM6.4 were shown to form nanodomains when transiently overexpressed in *Nicotiana benthamiana* (Bücherl et al., 2017; Jarsch et al., 2014), indicating that REM C-terminal-domain-mediated homo-oligomerization is not strictly for nanodomain formation. These nanodomains may instead consist of REM dimers and/or REM hetero-oligomers. It is also possible that some REM oligomerize by mechanisms that AlphaFold3 cannot predict such as through interactions mediated by their intrinsically disordered domain (Marín & Ott, 2012).

The structural properties of REM C-terminal domains suggest that REMs may sense or impose nanoscale membrane curvature. Local curvature changes can influence protein and lipid lateral organization within membrane, as shown for instance for *Bacillus subtilis* chemoreceptors (Strahl et al., 2015). REM C-terminal domain curvature (Fig. 2 and Fig. 3), combined with their bar code-like positive surface charge (Xu et al., 2024), may confer lipid-binding selectivity and help define local membrane lipid composition. Changes in membrane curvature may also regulate mechanosensitive proteins, including mechanosensitive channels. Several REM members have been shown to associate with cytoskeletal regulatory components (Jolivet et al., 2023; Ma et al., 2022; Su et al., 2023) (von Arx et al., 2025), including membrane-spanning, cell-wall-associated, and actin-nucleating Formin 6 (Ma et al., 2022)( von Arx et al., 2025). Analogous to the mammalian integrin complex (Luciano et al., 2024) REMs may participate in plant cell mechanobiology, potentially explaining their role in salt and osmotic stress responses (Rui et al., 2025) (von Arx et al., 2025).

In addition to REM C-terminal domains intrinsic structural features, several factors are likely to influence how REMs sense and/or modify the plasma membrane. S-acylation of REM C-terminal anchors has been both predicted (Gronnier et al., 2017; Konrad et al., 2014) and experimentally confirmed (Fu et al., 2018; Hemsley et al., 2013; Konrad et al., 2014; Kumar et al., 2022). Such lipid modifications likely add another regulatory layer influencing membrane properties and curvature.

Given REMs’ propensity to form oligomers *in vitro* (Bariola et al., 2004; Marín & Ott, 2012; Martinez et al., 2019) and their intrinsically disordered domains (Marín & Ott, 2012), REM nanodomains have long been hypothesized to represent membrane-associated molecular condensates (Jaillais & Ott, 2020). Biomolecular condensates can modulate membrane organization, properties and curvature (Dragwidge & Van Damme, 2023; Mangiarotti et al., 2023; Meese & Strader, 2025). Thus, putative condensate-like behaviors in REM assemblies may provide an additional mechanism shaping REM-membrane interactions.

Transient overexpression in *Nicotiana benthamiana* suggested that REMs form distinct, co-existing nanodomains (Jarsch et al., 2014). Our analyses further predict that these nanodomains may differ strongly in their local membrane curvature. REMs are unlikely to be unique in this regard, and for instance BAR-domain membrane-shaping proteins found across Eukaryotes (Allen et al., 2022) may co-exist within cells. We therefore envision that, alongside the great diversity in plasma membrane nanodomain composition (Jaillais et al., 2024), multiple nanoscale membrane curvature states coexist in plant plasma membrane. Our in-silico analyses provide a conceptual framework for future experimental investigation.

## Acknowledgement

We thank all members of the NanoSignaling Laboratory for fruitful discussions and comments on the manuscript. This research was supported by the Deutsche Forschungsgemeinschaft (DFG) grants A08-SFB1101 and B01-TRR356 to J.G. and by the Technical University of Munich.

## Methods

### Sequence alignment and phylogenetic analyses

REMORINs homologues were searched iteratively using the protein sequences of the *Arabidopsis thaliana* (At) REMORINs AtREM6.2, AtREM6.3, AtREM6.4, AtREM3.1 AtREM4.1, AtREM6.6, AtREM1.1, AtREM1.2 and *Marchantia polymorpha* (Mp) REMORINs MpREM1, MpREM2, MpREM3, and BLASP 2.11.0+ (Camacho, 2009) with default parameters and e-value threshold of 1e^−10^, against a database of 53 plant species via Phytozome 13 (Goodstein et al., 2012) collecting over 1100 REM sequences (Supplementary Data 1). When several splice variants were identified for a single locus, the longest variants were kept for subsequent analyses. REMORINs protein sequences were aligned using MUSCLE v3.8.1551 (Edgar, 2004a, 2004b), in Jalview Version 2 (Waterhouse et al., 2009). Subsequently, the positions in the multiple sequence alignment with more than 80% of gaps were removed using trimAl v1.5.rev0 (Capella-Gutiérrez et al., 2009). For full length REMORIN sequences, no trimming was performed. The best-fit empirical amino-acid exchange rate matrix to be employed on full length and trimmed alignments were defined using ModelFinder (Kalyaanamoorthy et al., 2017) according to Bayesian information criterion. Maximum likelihood analyses were conducted using IQ-TREE3.0.1(Wong et al., 2025). Branch supports were assessed using 2 000 replicates of UltraFast Bootstraps (Hoang et al., 2018). The resulting consensus trees were visualized and analyzed using iTOL v7 (Letunic & Bork, 2024). To obtain the site-specific conservation score, evolutionary rates were extracted with IQ-TREE3.0.1 using an empirical Bayesian method (Wong et al., 2025) and scaled from 1-10. Sequences whose length exceeded 90% of all other sequences in each clade were removed from the analysis.

### Structure modelling and analyses

Proteins and protein complexes structures were predicted using AlphaFold3 (Abramson et al., 2024) and analyzed using ChimeraX (Pettersen et al., 2021). The analysis of amino acids conservation on REM1.2 C-terminal domain structure was performed using Consurf 2016 (Ashkenazy et al., 2016) using a full-length REM multiple sequence alignment (described above). Position-specific evolutionary rates were calculated under an empirical Bayesian methodology and normalized and grouped into nine conservation grades, where one corresponds to most rapidly evolving positions and nine correspond to the most evolutionarily conserved positions. The analyses of REM C-terminal domain dimers lengths and curvatures were performed using custom Python scripts (https://github.com/NanoSignalingLab/Biermann_002.git) and Python 3.11, NumPy, SciPy, Matplotlib, and Pandas. Cα coordinates were extracted from PDB files and projected onto the XY plane. A circle was fitted using nonlinear least-squares minimization. For each atom, the radial distance from a candidate circle center (x_c_,y_c_) was calculated as:

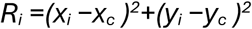

The optimal circle center was defined by minimizing the deviation of *R_i_* from their mean value. The fitted radius (*R*) was taken as the mean radial distance after optimization, and the two-dimensional curvature (*κ*) was defined as:

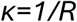

The putative membrane-binding interface was estimated as the arc length of the fitted circle spanned by the Cα atoms, calculated as:

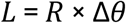

where *L* is the interface length (Å), *R* is the circle radius (Å), and Δθ is the angular span in radians.

To evaluate local structural bending, a B-spline curve was fitted through the ordered Cα coordinates. The first and second derivatives of the spline were computed, and local curvature was defined as:

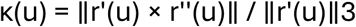

where r(u) is the spline parameterization of the backbone, r′(u) its first derivative, and r′′(u) its second derivative. The mean spline curvature was calculated as the average of κ(u) across the entire curve, while the maximum spline curvature corresponds to the largest local value. All analyses were automated in batch mode, and diagnostic plots (circle fit, interface arc, angle histogram) were generated for manual quality control.

---

**Figure S1.**
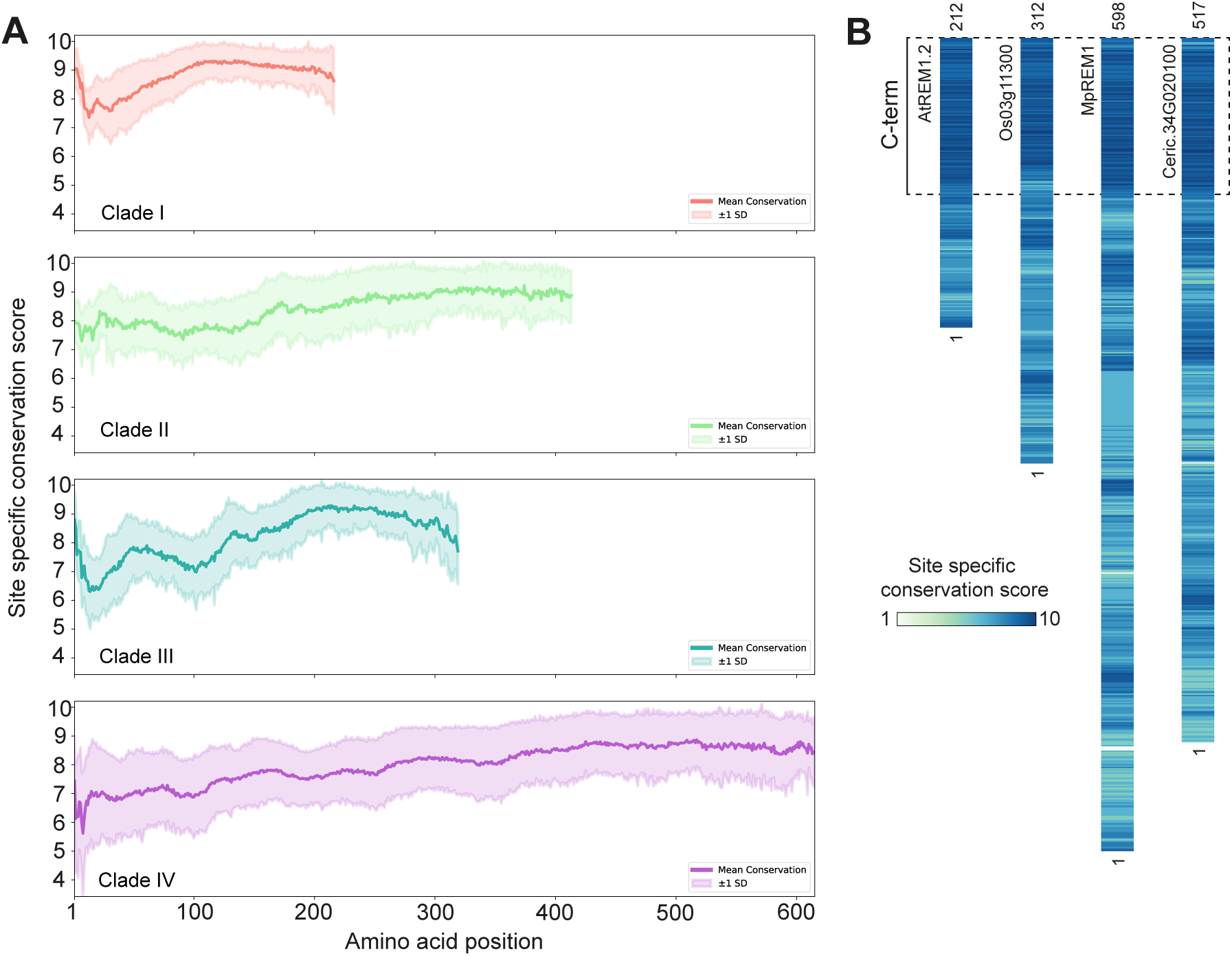
Analysis of site-specific amino acid conservation. Site-specific evolutionary rates were calculated under an empirical Bayesian methodology, normalized and grouped into ten conservation grades, where one corresponds to most rapidly evolving positions and nine correspond to the most evolutionarily conserved positions. Site-specific evolutionary rates are shown as the average conservation per clade in **A** and shown for one REM from each clade in **B**.

**Figure S2.**
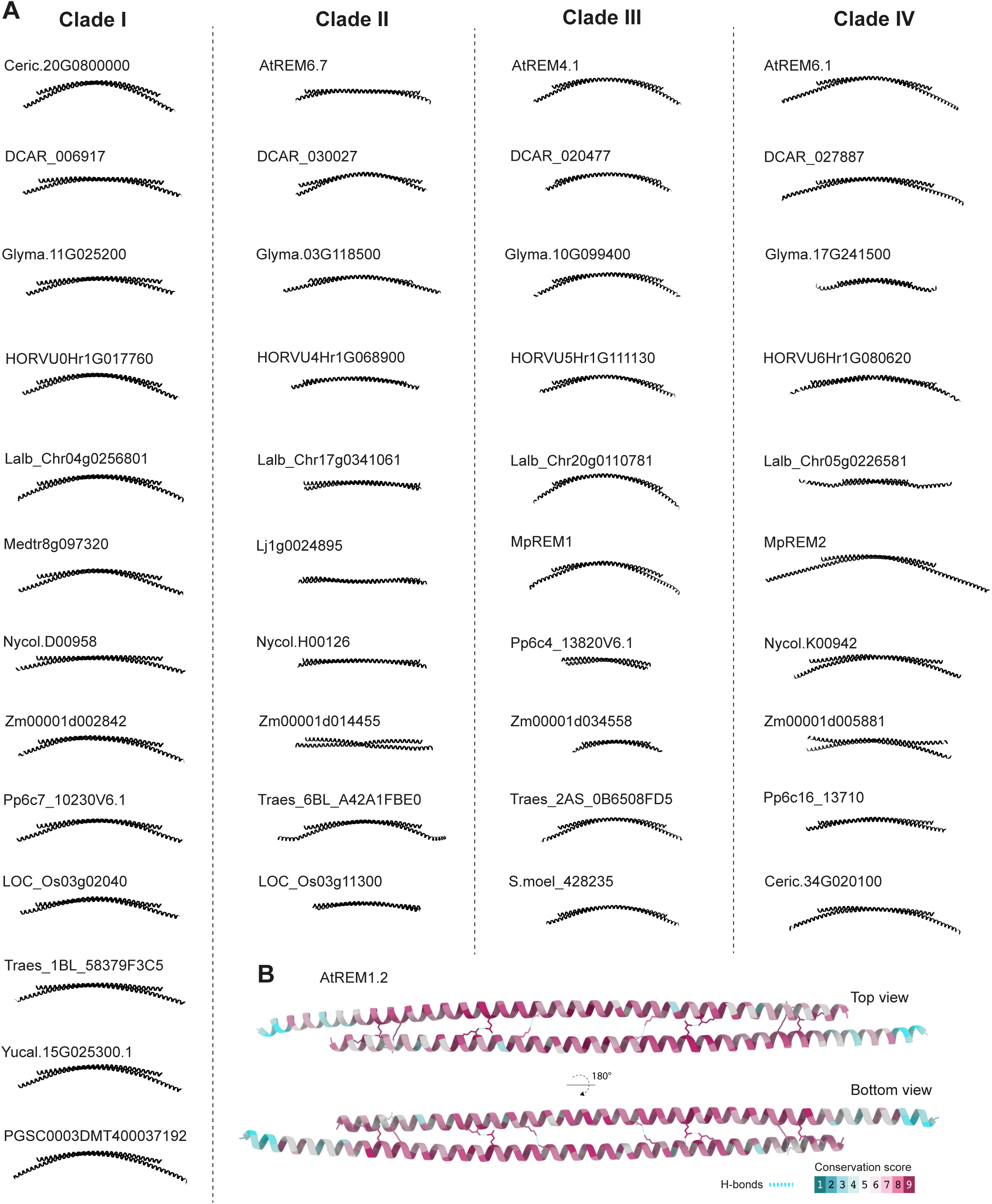
AlphaFold3 analyses of REM dimers. **A**. AlphaFold3 predicted structure of REM C-terminal domain dimers for 43 REMs. **B**. Analysis of the amino acid conservation across 1133 REMs from four clades. Amino acids engaged in dimer-dimer interaction are strongly conserved.

**Figure S3.**
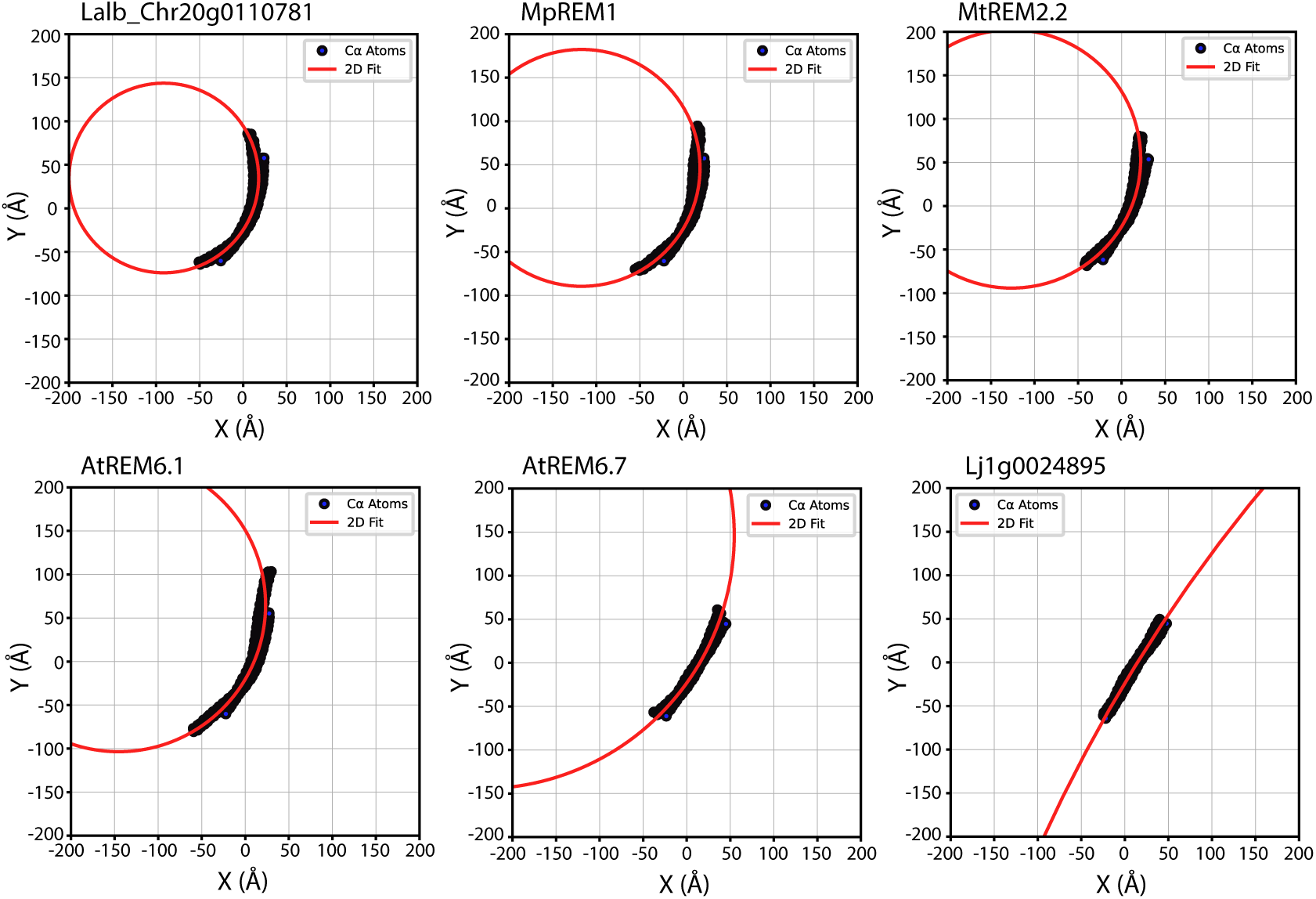
Estimation of REM membrane-binding interface features. Examples of nonlinear least-squares minimization circle fitting to the Cα atoms backbone of AlphaFold3 predicted structures of the *Lupinus albus* Lalb_Chr20g0110781, *Marchantia polymoprha* MpREM1, the *Medicago truncatula* MtREM2.2, the *Arabidopsis thaliana* AtREM6.1 and AtREM6.7, and *Lotus japanicus* Lj1g0024895 REMORIN dimers.

**Figure S4.**
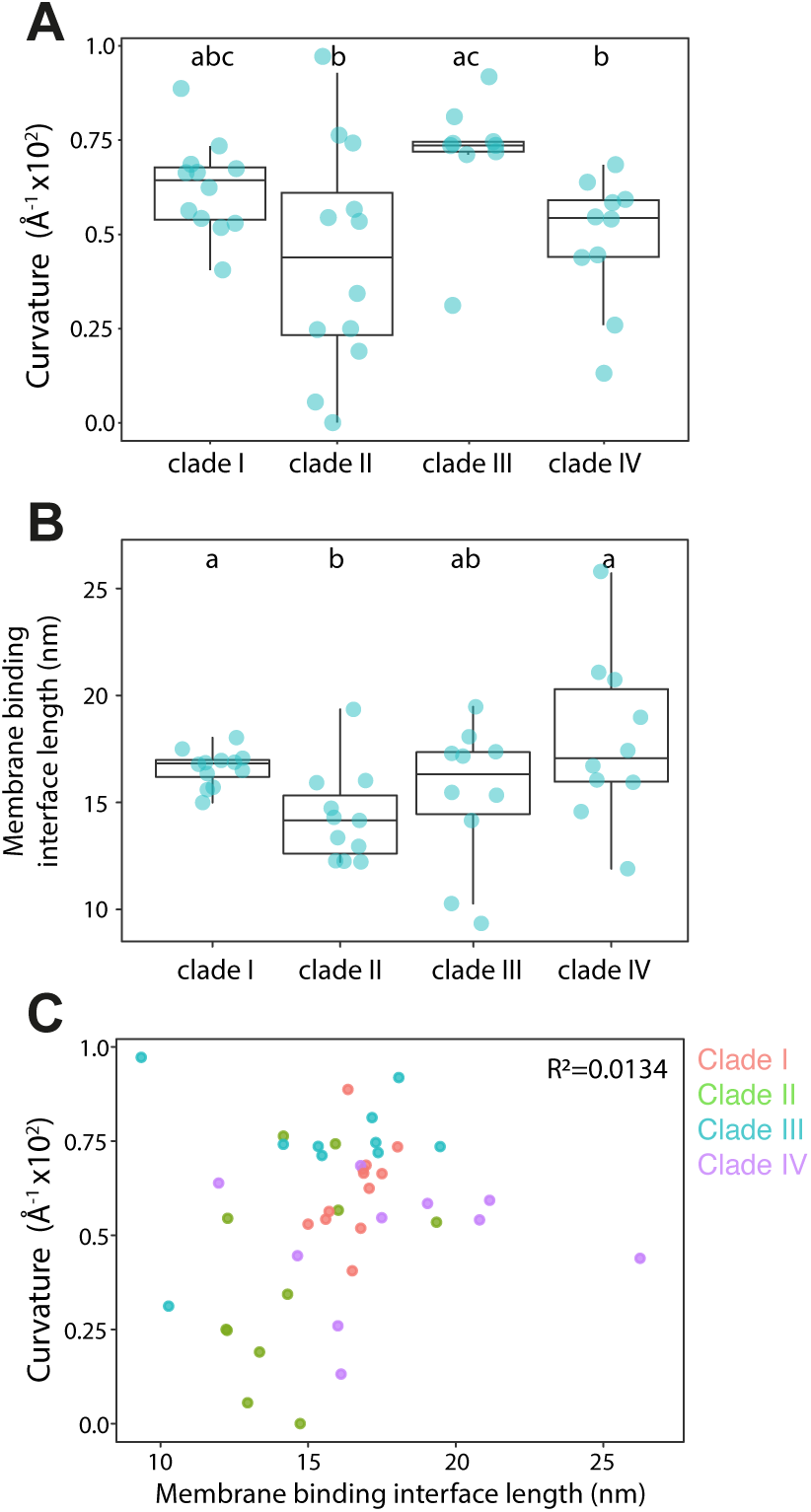
Analysis of the variation of the estimated curvatures and lengths of REM C-terminal membrane binding interface between REM clades. **A**. Curvatures of REM C-terminal domain dimers estimated using from nonlinear least-squares minimization circle fitting to AlphaFold3 predicted structures for each REM clade. Each datapoint presents the estimated curvature for one REM dimer. Conditions which do not share a letter are significantly different in Dunn’s multiple comparison test (p < 0.05). **B**. Lengths of REM membrane-binding interface estimated from the arc length of the fitted circle spanned by REM dimer Cα atoms for each REM clade. Each datapoint presents the estimated curvature for one REM dimer. Conditions which do not share a letter are significantly different in Dunn’s multiple comparison test (p < 0.05). **D**. Correlation analysis between estimated membrane binding interface length (nm) and estimated curvature (Å^-1^ x10^2^).

**Figure S5.**
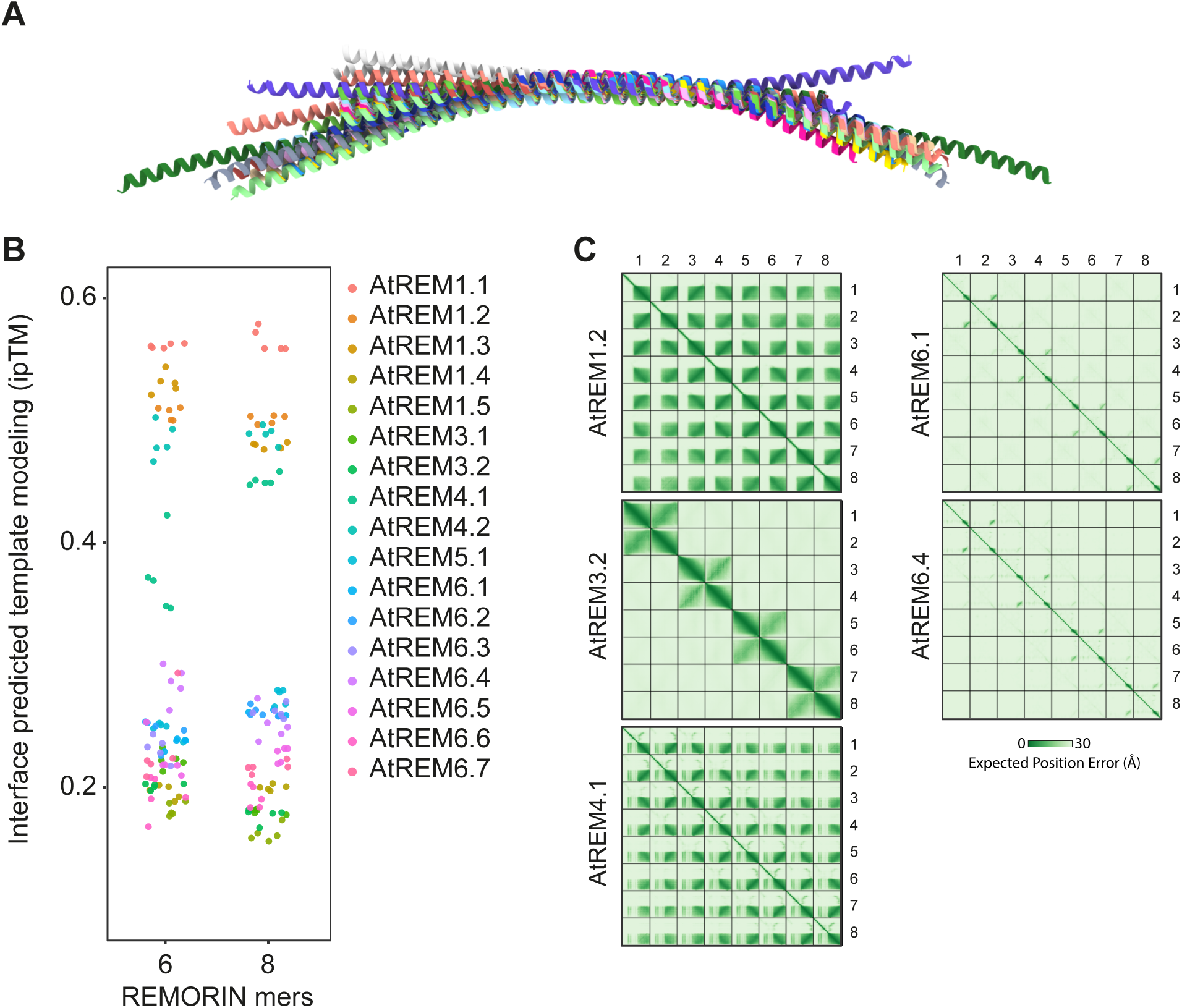
Analysis of the predicted oligomerization status of *Arabidopsis thaliana* REMs. **A.** Superimposed AlphaFold3 structure predictions of *Arabidopsis thaliana* REM C-terminal domain dimers. **B.** Interface predicted template modelling (ipTM) score of AlphaFold3 predictions for full-length *Arabidopsis thaliana* REMs hexamers and octamers. Each datapoint represents individual ipTM prediction scores for each multimer. **C.** Expected position error (Å) of AtREM1.2, AtREM3.2, AtREM4.1, AtREM6.1 and AtREM6.4 monomer chain with respect to the other chains in predicted multimers.

**Figure S6.**
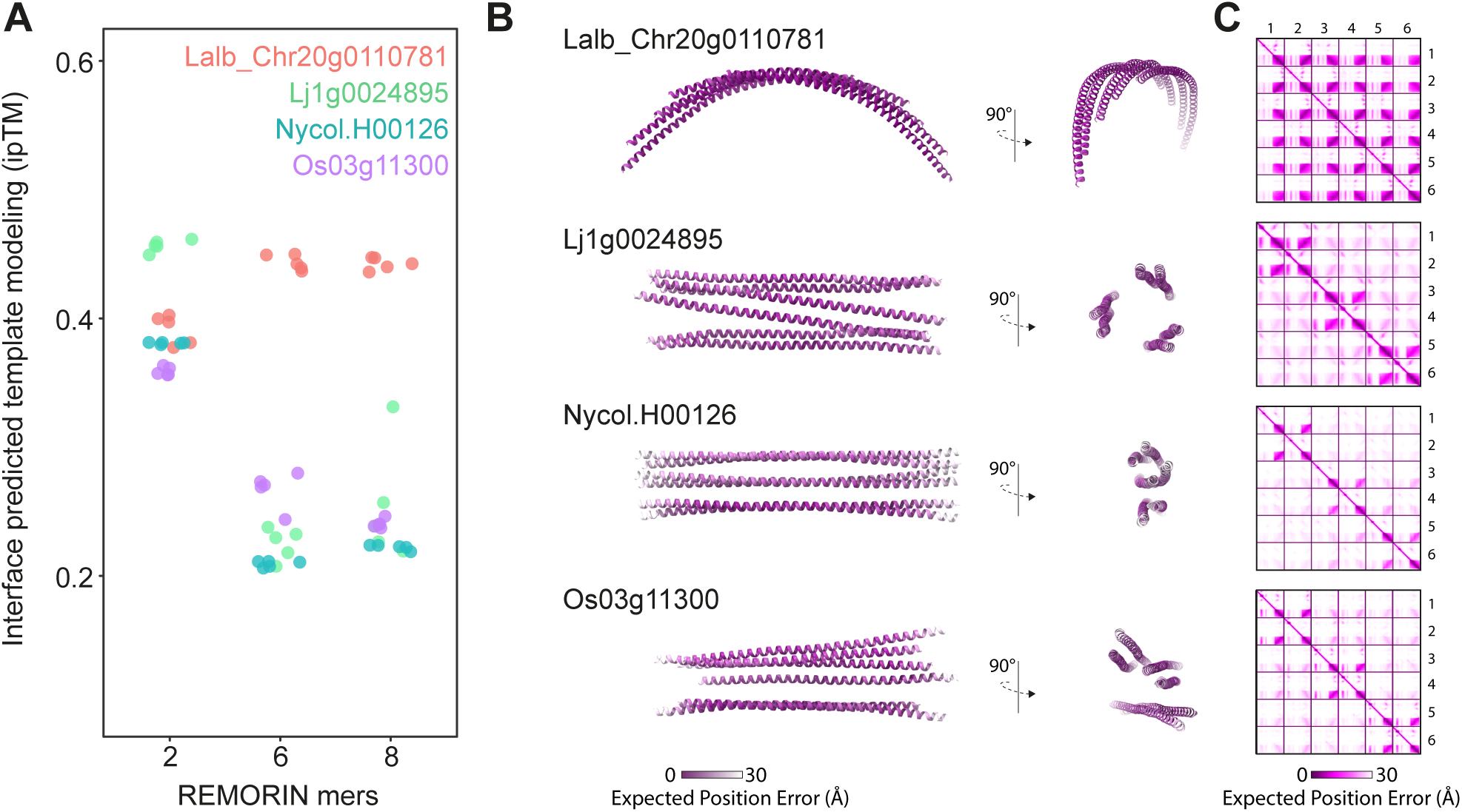
Analysis of the predicted oligomerization status of additional REMs. **A.** Interface predicted template modelling (ipTM) score of AlphaFold3 predictions for full-length REMs dimers, hexamers and octamers. Each datapoint represents individual ipTM prediction scores for each multimer. **B.** Corresponding AlphaFold3 predicted structures of the predicted oligomers. Structures are colored according to expected position error (Å) within single REM chains. **C.** Expected position error (Å) of Lalb_Chr20g0110781, Lj1g0024895, Nycol.H00126 and Os03g11300 monomer chain with respect to the other chains in predicted hexamers. Lalb_Chr20g0110781 is predicted to form hexamers while only dimeric assemblies can be predicted with confidence for Lj1g0024895, Nycol.H00126 and Os03g11300.

